# Integrin signaling downregulates filopodia in muscle-tendon attachment

**DOI:** 10.1101/270546

**Authors:** Benjamin Richier, Yoshiko Inoue, Ulrich Dobramysl, Jonathan Friedlander, Nicholas H. Brown, Jennifer L. Gallop

**Affiliations:** The Gurdon Institute, Tennis Court Rd, Cambridge CB2 1QN, United Kingdom; Dept. of Biochemistry, University of Cambridge; Dept. of Physiology, Development and Neuroscience, University of Cambridge

**Author notes:** Joint first authors (Benjamin Richier, Yoshiko Inoue). Correspondence to Jennifer Gallop.

**Keywords:** filopodia, integrin signaling, actin dynamics, muscle, Drosophila

## Abstract

Cells in developing tissues need to sense their environment for their accurate targeting to specific destinations. This occurs in developing muscles, which need to attach to their respective tendon cell before muscle contractions can begin. Elongating myotube tips form filopodia, which are presumed to have sensory roles, and are later suppressed upon building the attachment site. Here, we use live imaging and quantitative image analysis of lateral transverse (LT) myotubes in *Drosophila* to show that filopodia suppression occurs as a result of integrin signaling. Loss of the integrin subunits αPS2 and βPS increased filopodia number and length at stages when they are normally suppressed. Conversely, inducing integrin signaling, achieved by expression of constitutively dimerised βPS cytoplasmic domain (diβ), prematurely suppressed filopodia. We discovered that the integrin signal is transmitted through the ArfGAP and scaffolding protein Git (G-protein receptor coupled interacting protein) and its downstream kinase Pak (p21-activated kinase). Absence of these proteins causes profuse filopodia formation and prevents filopodial inhibition by diβ. Thus, integrin signaling switches off the exploratory behaviour of myotubes seeking tendons, enabling the actin machinery to focus on forming a strong attachment and assembling the contractile apparatus.

## Introduction

Muscle development involves similar phases of differentiation and movement in *Drosophila* and vertebrate embryos (Schweitzer et al., 2010). *Drosophila* somatic muscles display a highly stereotyped pattern of 30 muscles in each abdominal segment, near the surface of the embryo which means they can be easily identified and followed during live cell imaging (Figure 1A, A’). The lateral transverse (LT)muscles migrate and elongate towards the epidermis and their attachment sites are particularly accessible for imaging (Fig. 1A, A’). Each LT muscle is a single syncytial cell, the myotube, which forms by the fusion of fusion-competent myoblasts to a founder cell (Rochlin et al., 2010). Each founder cell contains the transcriptional instructions for the development of that particular muscle. This include regulating the number of fusion events and the sites of attachment when the resulting myotube polarizes and elongates bidirectionally to attach to the epidermis (Fig. 1B-B") (de Joussineau et al., 2012). While some of the guidance cues known to target developing myotubes to their tendon cell attachments overlap with those guiding neuronal cell migration (Slit, Robo and Derailed), some cues are unique to specific myotubes like the transmembrane protein Kon-tiki in ventral-longitudinal muscles (Callahan et al., 1996; Estrada et al., 2007; Kramer et al., 2001; Schnorrer et al., 2007). Myotube migration and tendon cell specification proceed in concert to organize muscle architecture (Schweitzer et al., 2010), with extensive protrusions occurring in both cell types at the time of attachment in the case of the adult indirect flight muscles (Weitkunat et al., 2014).

**Figure 1:**
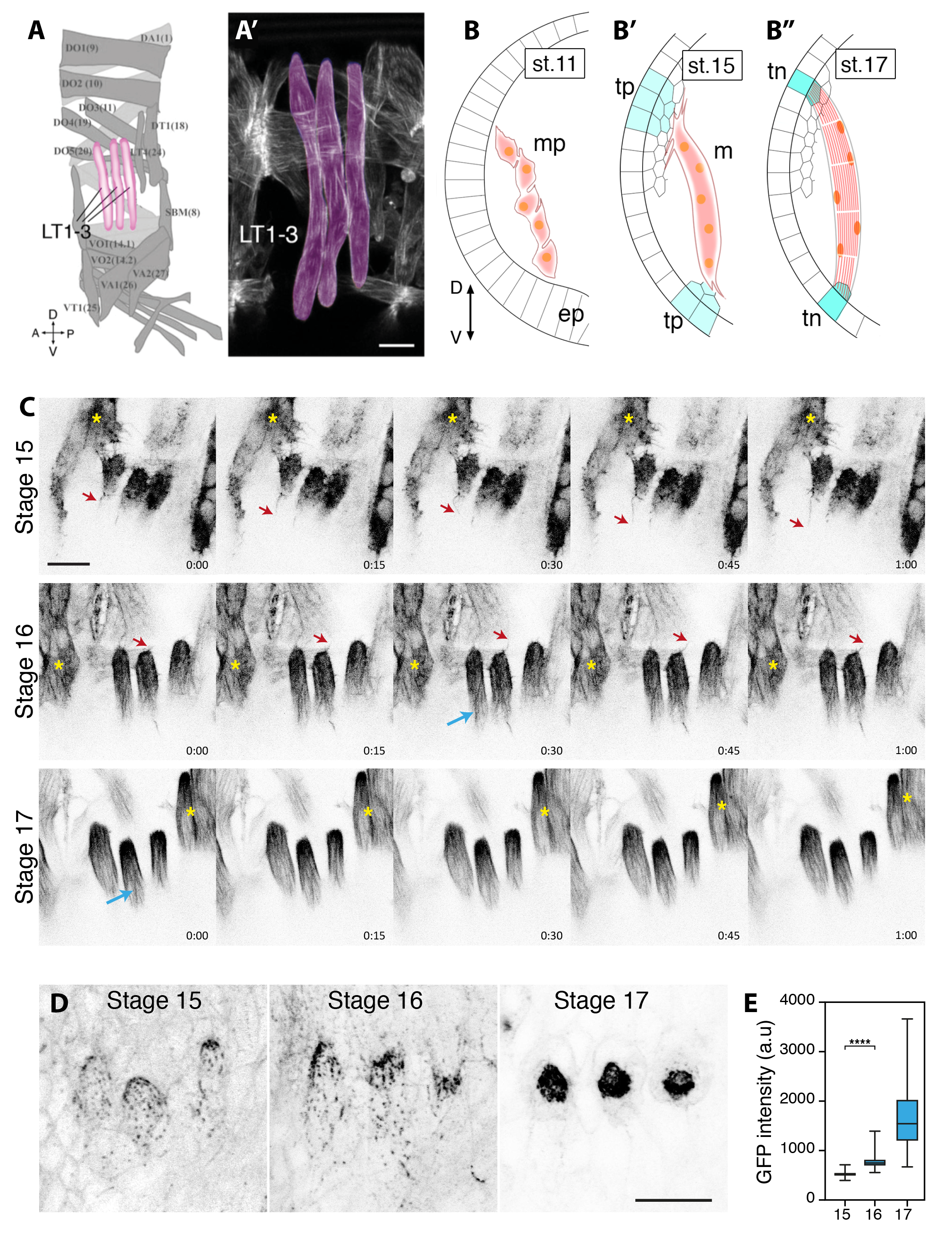
Filopodia dynamics in embryonic LT muscles. (A-A’) Schematic representation of the stereotyped muscle pattern found in segments A2 to A7 (Ruiz-Gomez et al., 1997). Lateral transverse muscles (LT1)3) are indicated magenta, interior muscles in light gray, other muscles in gray. Orientation: Dorsal (D) ventral (V), anterior (A) and posterior (P). (A’) Mature LT muscles (magenta) in a stage 17 Drosophila embryo visualized by GFP-actin under the control of mef2-Gal4. Scale bar = 10μm. (B-B”) Schematic representation of developmental steps of Drosophila myogenesis. Cross section of an embryo is shown, dorsal (D) up, ventral (V) down. (B) Phase 1 (stage 11): several fusion-competent myoblasts fuse to one founder cell to form a multinucleate myotube progenitor (mp). (B’) Phase 2 (stage 15): the leading edge of the extending myotubes form filopodia in search of their specific targets at the epidermis, the tendon precursors (tp) located in the epithelial layer (ep). (B’’) Phase 3 (stage 17): filopodia formation ceases, and both ends of the myotube attach to terminally differentiated tendon cells (tn). (C) Timelapse images of LT myotube tips visualized by GFP-actin expression driven by *mef2-Gal4* at stages 15, 16 and 17. Filopodia form during stage 15 and 16 but appear smaller at later stages (red arrows). Within the myotube, actomyosin fibers develop during stage 16 (blue arrows). Yellow asterisks indicate adjacent myotubes. Time = min:sec. Scale bar = 10 μm. (D-D”) Equivalent images of βPS-GFP expressed at endogenous levels showing integrin accumulation at muscle attachment site at stages 15, 16 and 17. Bar = 10 μm. (E) Quantification of βPSGFP intensity at myotube tips (8-10 embryos). Box and whiskers indicate the second and third quartile and the minimum and maximum values. The horizontal bar shows the median. ST15, n = 68; ST16, n = 85; ST17, n = 101. Mann-Whitney test: p = 2.2 × 10^−16^.

In the embryo, the myotube ends that move toward the tendon cells produce several actin-rich finger-like protrusions known as filopodia (Schnorrer and Dickson, 2004). Filopodia are implicated in the ability of cells to find their correct location (Davenport et al., 1993; Jacquemet et al., 2015). Once the right locations are reached, the myotubes need to stop migrating, otherwise secure attachments cannot form and myotubes ‘overshoot’ the tendon cells. Such a phenotype has been observed in *Drosophila* deficient in G protein-coupled receptor kinase interacting ArfGAP (Git) (Bahri et al., 2009). At tendon cells, myotubes make a strong integrin-based attachment, the protrusions reduce and the myotube tips become rounded. There is concerted assembly of the contractile actin machinery to form the sarcomeres, during which tension itself is implicated (Weitkunat et al., 2014). This developmental process illustrates the intracellular specificity in how the actin cytoskeleton is arranged,with the need for appropriate actin filament nucleators, elongators and bundlers to be recruited to different areas of the cell in a spatially and temporally coordinated manner. Remodeling of the actin cytoskeleton is proposed to occur actively, for example by the activation of Rho type GTPases or phosphorylation cascades. Within the context of myotube migration, one could envisage that the building of the extensive and specialized contractile machinery of the myotube could lead to a natural reduction in migratory, protrusive actin activity. Alternatively, the protrusive actin structures needed for migration could be actively regulated and suppressed at the appropriate time.

In this study, we investigate the role of integrins in the regulation of filopodia during the formation of the muscle-tendon attachment site. We find that, as the muscle reaches the tendon cell, integrins have two important activities: their well characterized role in attaching to the tendon cell via an intervening extracellular matrix, and a new function mediating a signaling pathway that actively suppresses filopodia. We characterize Git and its downstream target p21 activated kinase (Pak) as components of this signaling pathway. The transition from migration to a process of muscle-tendon attachment provides a rare example where the contribution of integrins to *Drosophila* development demonstrably involves signaling in addition to forming adhesions. Thus, integrin signaling helps to provide a migratory stop signal that acts in parallel with the formation of an integrin adhesion junction at the muscle ends.

## Results and Discussion

### Filopodia at the myotube leading edges are downregulated when the attachment site with the tendon cell is formed

Myotubes migrate towards prospective tendon cells in the epidermis and form stable attachment sites. Contractile actomyosin bundles then give rise to muscle contraction and embryo movement (Schweitzer et al., 2010) (Fig. 1A-B"). By observing expression of GFP-actin in the dorsal tip of LT muscles (using the muscle specific Gal4 driver mef2-Gal4) in live embryos, we visualized active filopodial dynamics (Fig. 1C, red arrows; Movies 1-3). Between stages 15 and 16 (~3 hours), dorsal LT tips went through morphological and behavioral changes. Dorsal LT tips were very close to each other with active filopodia dynamics during stage 15 and beginning of stage 16 (Fig. 1C, stage 15). As development progressed, LT tips became more spaced out and the number and length of filopodia reduced while actin bundles formed instead within the myotube (Fig. 1C, blue arrows), with a rounded end of the myotube making the attachment to the tendon cell (Fig. 1C, stage 16). At stage 17, filopodia had become very rare and muscle contractions had started (Fig. 1C, stage 17). This apparent simultaneous reduction of filopodia number and attachment formation is consistent with previous observations in embryonic ventral oblique muscles (Bate, 1990; Schnorrer and Dickson, 2004). The integrin expressed by the muscles is the heterodimer αPSβPS, which attaches the muscle to the tendon cell so that the force of muscle contractions is transmitted to the exoskeleton for movement (Maartens and Brown, 2015; Devenport et al., 2007). We observed,similarly to previous studies in indirect flight muscles development (Weitkunat et al., 2017),that the filopodial downregulation occurred at the same time as the accumulation of βPS integrin at the attachment sites (Fig. 1D, E).

### Integrin signaling downregulates filopodia

In order to test whether integrins are inhibiting filopodia formation, it was necessary to quantify and characterize the behavior of filopodia produced by muscles LT1-3 (as shown in Fig. 1A). To do this, we developed an image segmentation and analysis pipeline for the reconstruction of filopodia in timelapse movies of five confocal microscope Z-stacks. The location of the specific muscles of interest within a complex tissue environment that has other muscles nearby, and the fact that filopodia can extend in any direction, made the analysis too complex for automated filopodia quantification software. We therefore employed a semi-automated pipeline consisting of three main steps:(1) Automatic segmentation and extraction of line segment candidates belonging to filopodia by using the Frangi Vesselness measure (Frangi et al., 1998), which was developed to track blood vessels in MRI data, (2) manual editing to remove false detections and merging of segments belonging to the same filopodium, and (3) automated linking of edited filopodia across time points (see Methods section for more details). The pipeline outputs a list of filopodia and their segment coordinates for each time point of their existence. Because the reconstruction of very small filopodia was very noisy, we filtered our dataset to focus on filopodia that reached a maximum length of 3 μm or more and existed for at least two timeframes. From this data, we then extracted measures such as the maximum length distributions of filopodia and the average number of filopodia from the movies (a z projection is shown in Fig. 2A, Movie 1). We quantified the filopodia for the three muscles LT1-3 together, rather than individually, as the three myotubes are closely apposed and difficult to separate. Because we label all muscles, adjacent muscles (Fig. 2, yellow asterisks) can mask LT muscles tips or their filopodia(Fig. 2, red arrows) when generating Z projection images. The pipeline uses the original Z stacks, which allows us to prevent these issues during the reconstruction. Using this method, we validated that there is a strong reduction in filopodia number from stage 15 to stage 16 in control embryos (see ctrl in blue in Fig. 2D).

With this tool at hand, we visualized GFP-actin in embryos lacking βPS (*myospheroid*^*XG43*^, *mys*^*XG43*^). In *mys*^*XG43*^ mutant myotubes, the filopodia were resistant to the downregulation usually observed by stage 16 (Fig. 2A’, B’ and D; Movie 4). A similar effect of removing integrin function was observed in embryos lackingαPS2 (*inflated^B4^, if^B4^*) although further defects in muscle development in these mutants prevented us from making an accurate reconstruction of filopodia (Fig. S1 and Movie 5). *Inflated* is the only αPS subunit expressed in muscles, whereas αPS1 and αPS2 are expressed in tendon cells (Devenport et al., 2007). This suggests that integrin heterodimers are required within the muscles to regulate filopodia at myotube tips. To distinguish between the signaling and adhesion functions of integrins we used diβ, a chimeric protein in which the intracellular domain of βPS is fused to the extracellular and transmembrane domains of a constitutively active form of the receptor Torso (Martin-Bermudo and Brown, 1999). This fusion protein constitutively dimerizes the cytoplasmic domain of the βPS subunit, and mimics integrin signaling, even in the absence of endogenous integrin-mediated adhesion to the extracellular matrix (Martin-Bermudo and Brown, 1999). The expression of diβ via the mef2-Gal4 driver prematurely reduced filopodia number in stage 15 embryos to a level similar to that seen at stage 16 in wildtype embryos (Fig. 2C and D; Movie 6).Expression of this construct only in muscle cells also provides an independent test of whether the effect of integrin on the suppression of filopodia is cell autonomous rather than due to contributions from tendon cells.

**Figure 2:**
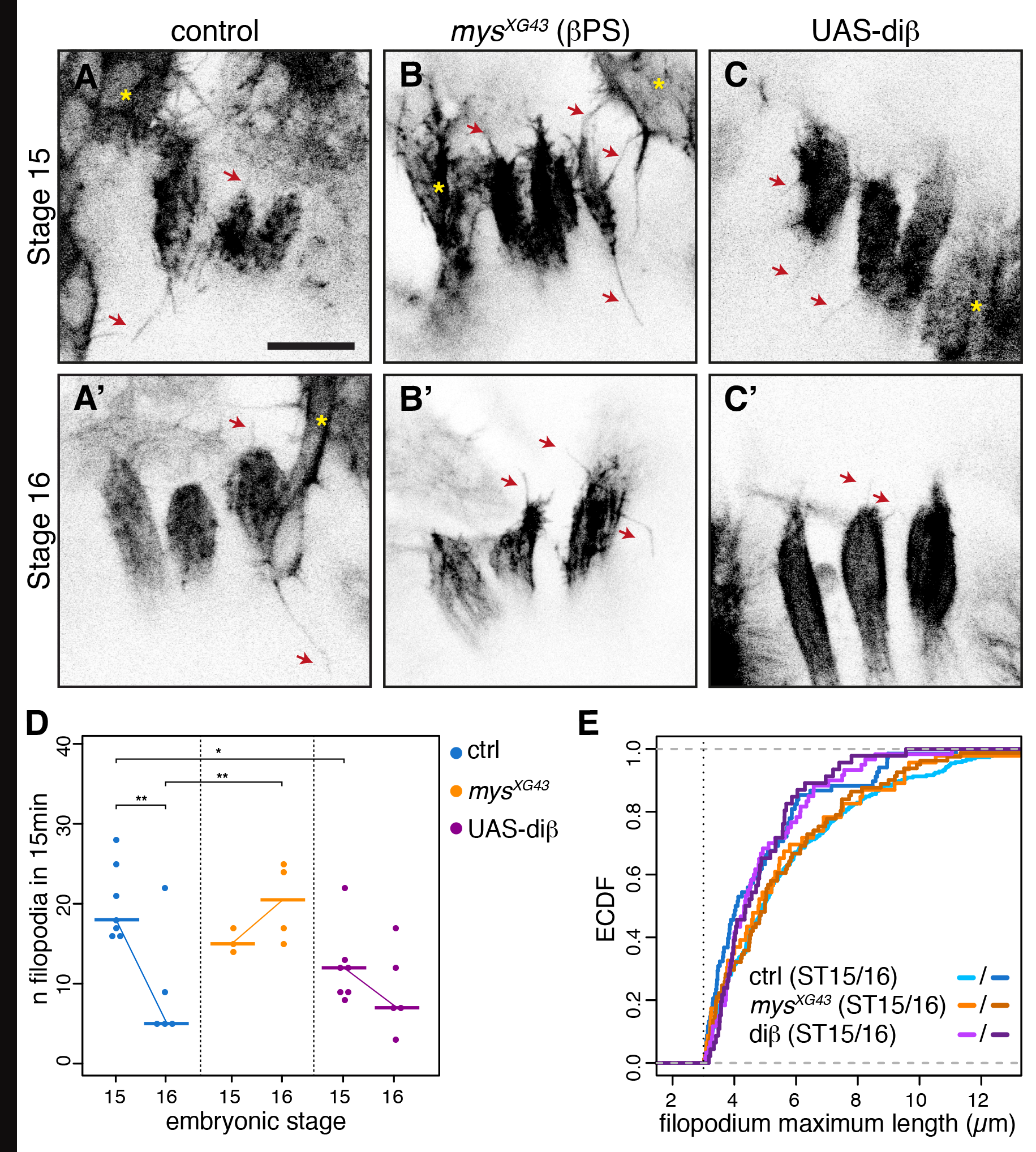
Integrins regulate filopodia formation at LT muscle tips. (A-C) Still sum intensity z projections from timelapse movies showing control(A-A’), zygotic βPS mutant (*mys*^*XG43*^ (B-B’) and UAS-diβ(C-C’) embryos at stage 15 and 16.Filopodia (red arrows) are visualized by actin-GFP expressed under control of *mef2-Gal4*.Yellow asterisks indicate adjacent muscles. All images are at the same scale. Scale bar: 10 μm. (D) Number of filopodia at LT tips during initiation and maturation of attachment site in a 15min time window. Individual values are represented by dots and horizontal bars indicate the median. Thin lines between stages illustrate the evolution of filopodia number. Non parametric Kruskal Wallis test, p=0.008753. Post hoc conover test,ctrl_st15 ~ ctrl_st16: p = 0.0001; ctrl_st16 ~ mys^XG43^_st16: p = 0.0014; ctrl_st15 ~ UAS-diβ_st15: p = 0.0116. (E) Because a large proportion of filopodia have a small maximum (max.) length in all conditions, we show here an empirical cumulative distribution function (ECDF) of the max. length values. A cut off at 3 μm was applied to discardthe shortest filopodia. Each point on the curve indicates the proportion of filopodia with a given max. length or less within the population. Thecurve for ctrl16 shifts towards the left compared to ctrl15, indicating that the proportion of long filopodia is smaller at stage16. Non parametricKruskal Wallis: p = 0.00111. *Post hoc* conover test (reject null hypothesis if p < α/2): ctrl_st15 ~ ctrl_st16: p = 0.0016; ctrl_st16 ~ mys^XG43^_st16: p = 0.0145; ctrl_st15 ~ UAS-diβ_st15: p = 0.0027. Sample size: ctrl_st15 = 264, ctrl_st16 = 68; mys^XG43^_st15 = 46, mys^XG43^_st16 = 81; UAS-diβ_st15 = 85, UAS-diβ_st16 = 46.

The finding that filopodia number was affected by both integrin loss and gain of function suggests an active role for integrin signaling in filopodia formation. To get further insights into this mechanism, we also analyzed the maximum length reached by each filopodia in our dataset. Maximum lengths of LT muscle filopodia follow an approximately exponential distribution (a high number of small filopodia and gradually fewer longer ones) (Fig. S2). This distribution is indicative of stochastic incorporation of actin generating filopodial length rather than a preferred length that is regulated by the cell. We display this as a normalized cumulative frequency chart (Fig. 2E) to make the differences between conditions easily visible. Similar to the reduction in numbers of filopodia, the lengths of the filopodia that are formed were reduced as the embryo develops from stage 15 to stage 16 (Fig. 2E, the line shifts to the left as there are proportionally fewer long filopodia). The filopodia in *mys*^*XG43*^ mutants at stage 15 were similar to controls, but at stage 16 the reduction in length of filopodia failed to occur (the stage 16 *mys*^*XG43*^ line overlays with stage 15 controls). Conversely, when diβ was expressed in the muscles, the maximum lengths distribution of stage 15 is shifted to the left and overlap stage 16 control (Fig. 2E). This shows that dominant active integrin signaling, by expression of diβ in the muscles, led to a premature reduction of filopodia maximum length compared to control embryos. The measure of filopodia length provides an independent quantification from filopodia number, hinting that integrin signaling may be affecting multiple points of actin regulation, affecting both the number and morphology of the resulting filopodia.

### Git and Pak control filopodia formation downstream of integrins

We next investigated the signals and pathways downstream of integrins in LT muscles during attachment to the tendon. G protein-coupled receptor kinase inteacting ArfGAP (Git) protein has been implicated in the integration and transduction of various signals at the membrane, including integrins and receptor tyrosine kinases (Hoefen and Berk, 2006). Git recruits Pak to sites of focal adhesions, as part of its complex with PIX (Pak-interacting exchange factor) and at muscle ends (Bahri et al., 2009; Hoefen and Berk, 2006). Moreover, *Git* mutants have previously been reported to have defects in muscle patterning and targeting in ventral muscles (Bahri et al., 2009). In these mutants, the muscles target and attach inappropriately, suggesting a defect in myotube tips sensing tendon cell sites of attachment.

In embryos lacking both maternal and zygotic contributions of Git (*Git* mz), LT myotubes displayed a more elongated and pointed tip instead of the rounded shape observed in wild type embryos both at stage 15 and 16 (Fig. 3A-B’). The quantification of filopodia indicates that the normal reduction in filopodia number at stage 16 failed to occur (Fig. 3F). As with loss of integrin, the filopodia appeared longer than in controls, but in this case also appeared more branched (Fig. 3B-B’, blue arrows). In addition, large protrusions, similar to lamellipodia, formed at muscle ends (Fig. 3B-B’, red arrowheads; Movie 7), which unfortunately prohibited an accurate quantification of filopodia length. A very similar phenotype was observed in embryos lacking both maternal and zygotic contributions of Pak (*Pak* mz) (Fig. 3C-C’; Movie 8). Thus, Git and its downstream effector Pak are also required to suppress filopodia at stage 16 and have an additional role in regulating the morphology of actin protrusions. The appearance of these larger protrusions when muscles lack Git or Pak suggests that there is normally a basal level of signaling through Git to suppress these structures. This is consistent with defects caused by the absence of Git in ventral muscles prior to the process of attachment, at Stage 14-15 (Bahri et al., 2009).

**Figure 3:**
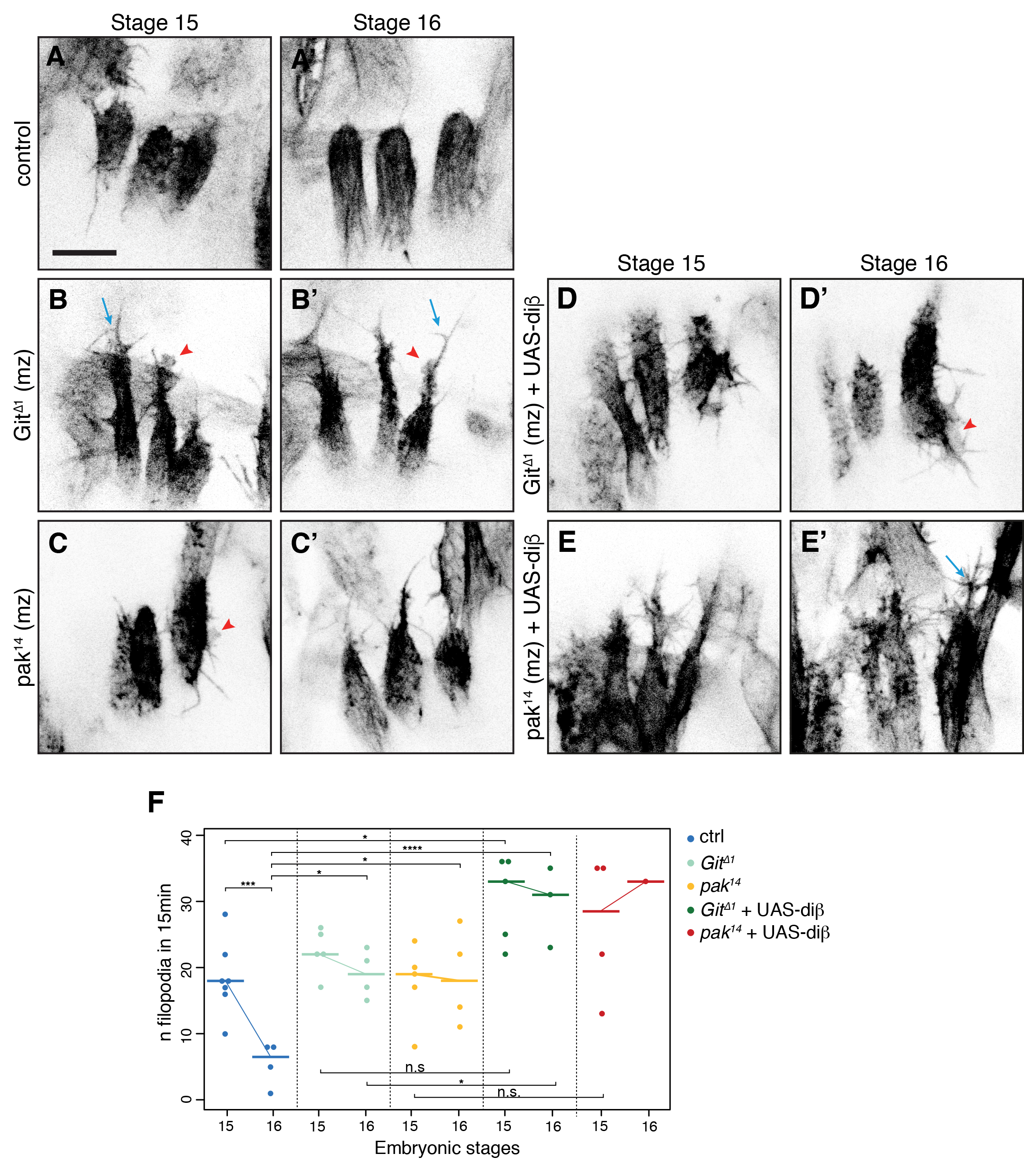
Integrins act upstream of Git and pak to suppress filopodia activity. (A-E’) Example images from timelapse movies for control (A-A’), *Git*^*Δ1*^ (B-B’), *pak*^*14*^ (C-C’) an UAS-diβ( expression in a *Git*^*Δ1*^ mz (D-D’) and *pak*^*14*^ mz (E-E’) background. Blue arrows show branched filopodia and red arrowheads indicate large lamellipodia-like structures. Yellow asterisks indicate adjacent myotubes. All images are at the same scale. Scale bar: 10 *μ*m. (F) Quantification of filopodia number at dorsal LT muscle tips. Horizontal bars show the median. Non parametric Kruskal wallis test, p = 0.00561. *Post hoc* conover test (reject null hypothesis if p < α/2), ctrl_st15 ~ ctrl_st16: p = 0.0080; ctrl_st16 ~ Git^Δ1^_st16: p = 0.0132; ctrl_st16 ~ pak^14^_st16: p = 0.0115; ctrl_st15 ~ Git^Δ1^ + diβ_st15: p = 0.0146; ctrl_st16 ~ Git^Δ1^ + diβ_st16: p = 0.00046; ctrl_st16 ~ pak^14^ + diβ_st16: p = 0.00115; Git^Δ1^_st15 ~ Git^Δ1^ + diβ_st15: p = 0.045; Git^Δ1^_st16 ~ Git^Δ1^ + diβ_st16: p = 0.014; pak^14^_st15 ~ pak^14^ + diβ_st15: p = 0.045; pak^14^_st16 ~ pak^14^ + diβ_st16: NA.

To test whether Git and Pak act downstream of integrins, or via parallel pathways, we expressed diβ in embryos lacking Git or Pak. The ability of diβ to prematurely suppress filopodia at stage 15 was lost in the absence of Git or Pak (Fig. 3D-F; Movies 9-10). The number of filopodia in embryos deficient in Git or Pak and expressing diβ were somewhat elevated further than either *Git* or *Pak* alone (however only one of the comparisons reached statistical significance). If true, this maybe an indication of an interplay between integrin signaling and adhesion. Dominant-negative effects of diβ on adhesion have been reported(Tanentzapf et al., 2006), and may serve to compound the protrusive defects observed in the absence of Git and Pak. Further studies will be required to investigate the connection between integrin signaling and adhesion in normal conditions. Overall, we conclude that Git and Pak act downstream of integrin signaling to suppress filopodia.

## Conclusions

During morphogenesis, cells must simultaneously integrate several signals and adapt their morphology and behavior to the developmental steps they go through, in coordination with their environment. During their attachment to tendon cells, myotubes need to recognize their target partner, stop elongating, start forming an adhesion and prepare the actomyosin structures that will produce muscle contractions, all in a short time period. In this study, we show that filopodia, which can be used to explore and identify tendon cells and are relevant during myotube elongation, are actively suppressed during muscle attachment. Integrins, which accumulate at the myotube tip to form a strong adhesion with tendon cells, also provide a signal in parallel acting through Git and Pak proteins. This is needed for the regulation of filopodia formation in order to ensure correct myotube arrest and adhesion to tendon cells.

It is likely that there are multiple proteins within the cytoskeletal machinery that are targeted by Git and Pak, possibly by direct phosphorylation or indirect regulation. Ena, the Scar/WAVE complex and fascin are all known actin regulators sensitive to phosphorylation state and present possible candidates downstream of Git and/or Pak (Comer et al., 1998; Loureiro et al., 2002; Mendoza, 2013; Zanet et al., 2012; Zeng et al., 2017). Further studies will be required to understand the intricate interactions between integrin adhesion and signaling and the actin cytoskeleton that allow the fine regulation of cell shape and behavior. To conclude, our results demonstrate that filopodia suppression occurs via integrin signaling within the myotube, providing a valuable new paradigm for integrin signaling in *Drosophila* and the roles of integrins in filopodia regulation.

## Acknowledgements

We thank A. Sossick and the Gurdon Institute imaging facility for microscopy equipment and assistance. We thank Jonathan Gadsby and Bishara Marzook for helpful comments on the manuscript. This work was supported by Wellcome Trust grant 069943 and 086451 to NHB, European Research Council Grant 281971 to JLG and Wellcome Trust Research Career Development Fellowship WT095829AIA to JLG. We acknowledge the core funding provided by the Wellcome Trust (092096) and CRUK (C6946/A14492). U.D. was supported by a Junior Interdisciplinary Fellowship via Wellcome Trust grant No. 105602/Z/14/Z and a Herchel Smith Postdoctoral Fellowship.

## Materials and Methods

### *Drosophila* stocks

Flies were raised and crossed at room temperature. The wild-type strain used was *white[1118]*. *Mef2-Gal4* was used to drive expression of these UAS constructs: *w*; P{UAS-pGFP.Act88F}1-2* (BL9254); *w*; P{UAS-pGFP.Act5C}2-1* (BL 9258) all from the Bloomington stock center, and UAS-diβ (Martin-Bermudo and Brown, 1999). The genomic rescue constructs used in this study was: *βPS-GFP* (Klapholz et al., 2015). The mutant lines include: *mys[XG43] (Bunch et al., 1992), if[B4]* (Brown, 1994), *Git[Δ1]* (in this study) and *pak[14]* (Newsome et al., 2000). The *FRT82B pak[14]* stock was ordered from the Bloomington stock center, *Git[Δ1]* and *pak[14]* germline clones were generated using *FRTG13 Git[Δ1]* and *FRT82B pak[14]* flies individually, essentially as described (Chou and Perrimon, 1996).

### Mutagenesis

The fly line carrying the P-element insertion *P{EPgy2}EY09254* was used to isolate mutations in *Git*. Mobilization of the P-element was done using the immobile element *P[ry+ Δ2-3](99B)* as a transposase source (Robertson et al., 1988). The genotype of the transposase stock is *Sp/CyO; P[ry+ Δ2-3](99B)/TM6*. Single jump starter males of the genotype *P{EPgy2}EY09254/CyO; P[ry+ Δ2-3](99B)/+* were crossed to second chromosome balancer females. From each cross, three white-eyed male progeny were crossed individually to a deficiency *Df(2R)Stan2, FRT42D/CyO* to screen for lethality. The *Git* alleles were analyzed by PCR for mapping of lesions in the *Git* locus. The excision allele *Git[Δ1]* deletes 1472 bp including the ATG and the first exon (−520 to 952 corresponding to the first ATG of the *Git* gene).

### Live imaging of *Drosophila* embryos

Dechorionated embryos (washed in 50% bleach) were mounted on a glass-bottomed dish with heptane glue and covered with water. In some cases, the genotype of embryos was determined by the absence of YFP-marked chromosome balancer prior and after the recording. Live imaging was performed on an inverted Leica TCS-SP5 equipped with a 63x 1.4 NA Plan Apo oil immersion objective or a Olympus FV-1000 confocal laser scanning microscope equipped with a 60x 1.3 NA Plan Apo oil immersion lens at room temperature. To follow filopodia movement we visualized stage 15-17 embryos in the indicated genetic backgrounds. Timelapse of LT tips in embryos were performed by taking z-stacks of 5-7 z sections (0.7 *μ*m spacing) at 15-second intervals for 15 min.

### Data analysis

The segmentation pipeline takes every Z-stack acquired throughout an imaging session separately. It outputs filopodia segments that are then edited by a user, followed by an automatic linking step that assigns filopodia identities across time points. The segmentation pipeline consists of the following seven steps:

1. Pixel segmentation: We use the *slic* segmentation algorithm (Achanta et al., 2012) without enforced connectivity to turn the greyscale frame into a black/white segmented frame without blurring. We allow three labels to give the algorithm enough finesse to resolve filopodia and then consider every pixel that the algorithm assigns a label to as foreground.
2. Cell body identification: In order to identify cell bodies to mask the cell interior, we employ aggressive gaussian blurring (σ=5) together with adaptive thresholding with a large (999×999 pixels) gaussian-weighed neighborhood (van der Walt et al., 2014) together with an erosion operation with a 5-pixel disk as stencil. We then use this as a mask to remove parts of the skeleton that lie inside cell bodies.
3. Vesselness filter: We employ a modified version of the Frangi Vesselness measure that is popular for tracing blood vessels in MRI data (Frangi et al., 1998). This filter employs the eigenvalues of the Hessian, which describe the local curvature of the image signal. In particular, the smallest of the eigenvalues describes the local“saddleness”. We therefore calculate the eigenvalues **λ**_1_, **λ**_2_ and **λ**_3_ of the Hessian for each pixel in the output of step 1 and order them such that |**λ**_1_|<|**λ**_2_|<|**λ**_3_|. We then calculate the derived quantities *r*_*a* =_|λ_2_|/|λ_3_|, 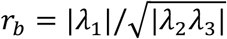 and 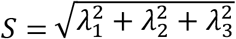. They are then combined into the Vesselness measure 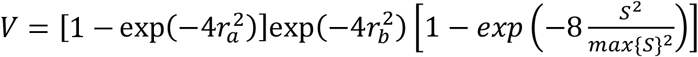. The exponential weights were chosen empirically such that filopodia in test movies were identified correctly.
4. Vesselness output segmentation: We again employ the slic segmentation algorithm without enforced connectivity, but a compactness parameter of 10 to identify bright connected lines in the Vesselness filter output and subsequently employ a binary closing step to heal broken lines.
5. Remove small objects: In this step, we remove small, connected regions containing less than 200 pixels (note that this step still operates on a Z-stack of images, hence the rather high pixel number threshold), which stem from noise and not actual filopodia.
6. Skeletonization: We use the 3D skeletonization algorithm by Lee et al. (Lee, 1994) to turn the resulting elongated patches into thin lines. We remove parts of the skeleton that are inside the cell bodies as detected in step 2.
7. Detection of connected line segments: We now label and iterate over all the connected regions of the skeleton. Each region has a collection of connected pixel coordinates (forming branched lines). We algorithmically split them into line segments and save them for subsequent manual editing and merging.

Following the automated extraction of filopodial line segments, we manually remove line segments that either were erroneously detected (due to e.g. noise in the image, or from parts of other structures in the field of view) or are part of filopodia stemming from cells other than the LT muscle cells. We also join line segments forming part of the same filopodium.

After the manual editing step, we automatically link and identify filopodia across time points. For each time point, we use the coordinates of all pixels contained in all filopodia over the preceding 5 time points. For each filopodium in the present time point, we then perform a next-neighbor search to find all previous filopodia that are close (with a cutoff radius of 25 pixels). For each of the candidate matches we calculate the path length difference measure *M*_*L*_ = |*L*_*c*_ - *L*_*p*_/*max*{*L*_*c*_,*L*_*p*_} (where L_c and L_p are the path lengths of the current filopodium and the candidate match filopodium, respectively) and the average direction measure 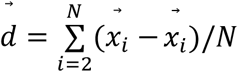(Where 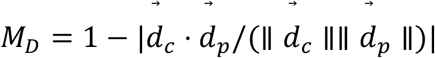 is the average non-normalized direction vector of a filopodium). We then summarize these measure into a score *S*= 0.46*M*_*L*_ + 0.6*M*_*D*_ (where the weights were manually adjusted to yield optimal results)and store the current filopodium and the match candidate together with the score. The pairs are then sorted by their score (lowest to highest) and iteratively linked until no more possible matches can be performed. Any leftover filopodia that werenșt matched to previously existing filopodia are considered to be newly created. After this step we save the list of filopodia and their segment coordinates in each frame in which they exist and extract quantities such as the maximum filopodia length (over the lifetime of a filopodium) and the number of filopodia in existence in each movie.

